# Counteracting estimation bias and social influence to improve the wisdom of crowds

**DOI:** 10.1101/288191

**Authors:** Albert B. Kao, Andrew M. Berdahl, Andrew T. Hartnett, Matthew J. Lutz, Joseph B. Bak-Coleman, Christos C. Ioannou, Xingli Giam, Iain D. Couzin

## Abstract

Aggregating multiple non-expert opinions into a collective estimate can improve accuracy across many contexts. However, two sources of error can diminish collective wisdom: individual estimation biases and information sharing between individuals. Here we measure individual biases and social influence rules in multiple experiments involving hundreds of individuals performing a classic numerosity estimation task. We first investigate how existing aggregation methods, such as calculating the arithmetic mean or the median, are influenced by these sources of error. We show that the mean tends to overestimate, and the median underestimate, the true value for a wide range of numerosities. Quantifying estimation bias, and mapping individual bias to collective bias, allows us to develop and validate three new aggregation measures that effectively counter sources of collective estimation error. In addition, we present results from a further experiment that quantifies the social influence rules that individuals employ when incorporating personal estimates with social information. We show that the corrected mean is remarkably robust to social influence, retaining high accuracy in the presence or absence of social influence, across numerosities, and across different methods for averaging social information. Utilizing knowledge of estimation biases and social influence rules may therefore be an inexpensive and general strategy to improve the wisdom of crowds.

## 1. Introduction

The proliferation of online social platforms has enabled the rapid expression of opinions on topics as diverse as the outcome of political elections, policy decisions, or the future performance of financial markets. Because non-experts contribute the majority of these opinions, they may be expected to have low predictive power. However, it has been shown empirically that by aggregating these non-expert opinions, usually by taking the arithmetic mean or the median of the set of estimates, the resulting ‘collective’ estimate can be highly accurate [1–6]. Experiments with non-human animals have demonstrated similar results [7–13], suggesting that aggregating diverse estimates can be a simple strategy for improving estimation accuracy across contexts and even species.

Theoretical explanations for this ‘wisdom of crowds’ typically invoke the law of large numbers [1, 14, 15]. If individual estimation errors are unbiased and center at the true value, then averaging the estimates of many individuals will increasingly converge on the true value. However, empirical studies of individual human decision-making readily contradict this theoretical assumption. A wide variety of cognitive and perceptual biases have been documented in which humans seemingly deviate from rational behavior [16–18]. Empirical ‘laws’ such as Stevens’ power law [19] have described the non-linear relationship between the subjective perception, and actual magnitude, of a physical stimulus. Such nonlinearities can lead to a systematic under-or over-estimation of a stimulus, as is frequently observed in numerosity estimation tasks [20–23]. Furthermore, the Weber-Fechner law [24] implies that log-normal, rather than normal, distributions of estimates are common. When such biased individual estimates are aggregated, the resulting collective estimate may also be biased, although the mapping between individual and collective biases is not well understood.

Sir Francis Galton was one of the first to consider the effect of biased opinions on the accuracy of collective estimates. He preferred the median over the arithmetic mean, arguing that the latter measure “give[s] a voting power to ‘cranks’ in proportion to their crankiness” [25]. However, if individuals are prone to under-or over-estimation in a particular task, then the median will also under-or over-estimate the true value. Other aggregation measures have been proposed to improve the accuracy of the collective estimate, such as the geometric mean [26], the average of the arithmetic mean and median [27], and the ‘trimmed mean’ (where the tails of a distribution of estimates are trimmed and then the arithmetic mean is calculated from the truncated distribution) [28]. Although these measures may empirically improve accuracy in some cases, they tend not to address directly the root cause of collective error (*i.e.*, estimation bias). Therefore, it is not well understood how they generalize to other contexts and how close they are to the optimal aggregation strategy.

Many (though not all) models of the wisdom of crowds also assume that opinions are generated independently of one another, which tends to maximize the information contained within the set of opinions [1, 14, 15]. But in real world contexts, it is more common for individuals to share information with, and influence, one another [26, 29]. In such cases, the individual estimates used to calculate a collective estimate will be correlated to some degree. Social influence can not only shrink the distribution of estimates [26] but may also systematically shift the distribution, depending on the rules that individuals follow when updating their personal estimate in response to available social information. For example, if individuals with extreme opinions are more resistant to social influence, then the distribution of estimates will tend to shift towards these opinions, leading to changes in the collective estimate as individuals share information with each other. In short, social influence may induce estimation bias, even if individuals in isolation are unbiased.

Quantifying how both individual estimation biases and social influence affect collective estimation is therefore crucial to optimizing, and understanding the limits of, the wisdom of crowds. Such an understanding would help to identify which of the existing aggregation measures should lead to the highest accuracy. It could also permit the design of novel aggregation measures that counteract these major sources of error, potentially improving both the accuracy and robustness of the wisdom of crowds beyond that allowed by existing measures.

Here, we collected five new datasets, and analyzed eight existing datasets from the literature, to characterize individual estimation bias in a well-known wisdom of crowds task, the ‘jellybean jar’ estimation problem. In this task, individuals in isolation simply estimate the number of objects (such as jellybeans, gumballs, or beads) in a jar [5, 6, 30, 31] (see Methods for details). We then performed an experiment manipulating social information to quantify the social influence rules that individuals use during this estimation task (Methods). We used these results to quantify the accuracy of a variety of aggregation measures, and identified new aggregation measures to improve collective accuracy in the presence of individual bias and social influence.

## 2. Methods

### 2.1. Numerosity estimation

For the five datasets that we collected, we recruited members of the community in Princeton, NJ, USA on April 26–28 and May 1, 2012, and in Santa Fe, NM, USA on October 17–20, 2016. Each participant was presented with one jar containing one of the following numbers of objects: 54 (*n* = 36), 139 (*n* = 51), 659 (*n* = 602), 5897 (*n* = 69), or 27852 (*n* = 54) (see Figure 1a for a representative photograph of the kind of object and jar used for the three smallest numerosities, and Figure S1 for a representative photograph of the kind of object and jar used for the largest two numerosities.). To motivate accurate estimates, the participants were informed that the estimate closest to the true value for each jar would earn a monetary prize. The participants then estimated the number of objects in the jar. No time limit was set, and participants were advised not to communicate with each other after completing the task.

**Figure 1:**
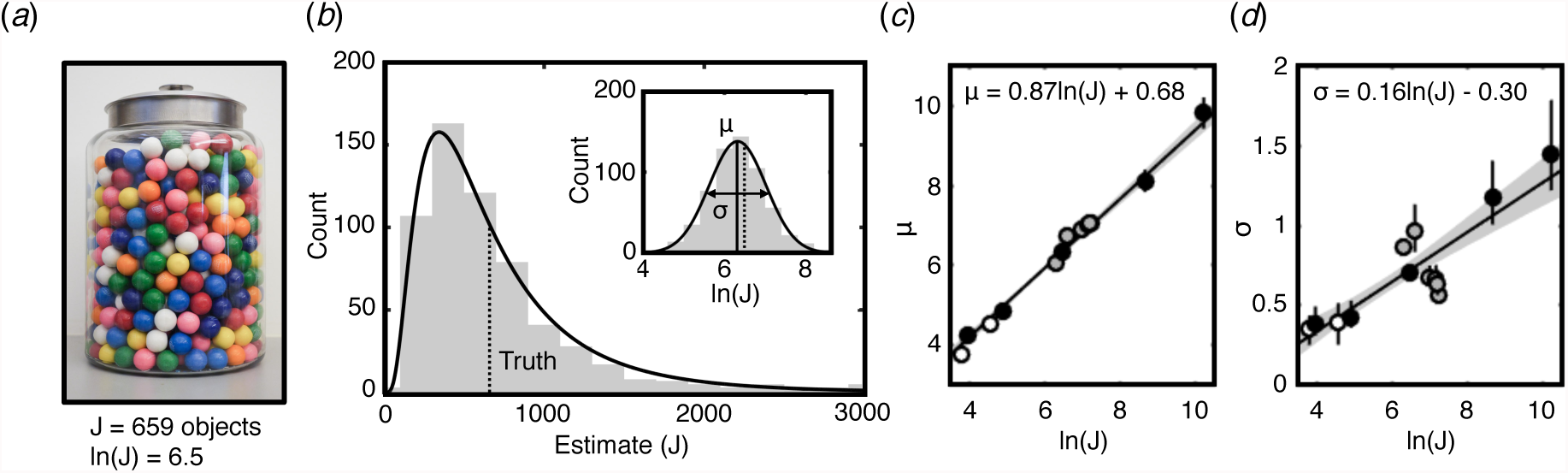
The effect of numerosity on the distribution of estimates. (a) An example jar containing 659 objects (ln(*J*) = 6.5). (b) The histogram of estimates (grey bars) resulting from the jar shown in (a) closely approximates a log-normal distribution (solid black line); dotted vertical line indicates the true number of objects. A log-normal distribution is described by two parameters, *µ* and *σ*, which are the mean and standard deviation, respectively, of the normal distribution that results when the logarithm of the estimates is taken (inset). (c-d) The two parameters *µ* and *σ* increase linearly with the logarithm of the true number of objects, ln(*J*). Solid lines: maximum-likelihood estimate, shaded area: 95% confidence interval. The maximum-likelihood estimate was calculated using only the five original datasets collected for this study (black circles); the eight other datasets collected from the literature are shown only for comparison (grey circles indicate other datasets for which the full dataset was available, white circles indicate datasets for which only summary statistics were available, see section 1 of the electronic supplementary material).

Eight additional datasets were included for comparative purposes and were obtained from refs. [5, 6, 30, 31]. Details of statistical analyses and simulations performed on the collected datasets are provided in the electronic supplementary material.

### 2.2. Social influence experiment

For the experiments run in Princeton (number of objects *J* = 659), we additionally tested the social influence rules that individuals use. The participants first recorded their initial estimate, *G*_1_. Next, participants were given ‘social’ information, in which they were told that *N* = {1, 2, 5, 10, 50, 100} previous participants’ estimates were randomly selected and that the ‘average’ of these guesses, *S*, was displayed on a computer screen. Unbeknownst to the participant, this social information was artificially generated by the computer, allowing us to control, and thus decouple, the perceived social group size and social distance relative to the participant’s initial guess. Half of the participants were randomly assigned to receive social information drawn from a uniform distribution from *G*_1_*/*2 to *G*_1_, and the other half received social information drawn from a uniform distribution from *G*_1_ to 2*G*_1_. Participants were then given the option to revise their initial guess by making a second estimate, *G*_2_, based on their personal estimate and the perceived social information that they were given. Participants were informed that only the second guess would count toward winning a monetary prize. We therefore controlled the social group size by varying *N* and controlled the social distance independently of the participant’s accuracy by choosing *S* from *G*_1_*/*2 to 2*G*_1_. Details of the social influence model and simulations performed on these data are provided in the electronic supplementary material.

### 2.3. Designing ‘corrected’ aggregation measures

For a log-normal distribution, the expected value of the mean is given by *X*_mean_ = exp (*µ* + *σ*^2^*/*2) and the expected value of the median is *X*_median_ = exp (*µ*), where *µ* and *σ* are the two parameters describing the distribution. Our empirical measurements of estimation bias resulted in the best-fit relationships *µ* = *m_µ_* ln(*J*) + *b_µ_* and *σ* = *m_σ_* ln(*J*) + *b_σ_* (Figure 1c-d). We replace *µ* and *σ* in the first two equations with the best-fit relationships, and then solve for *J*, which becomes our new, ‘corrected’, estimate of the true value. This results in a ‘corrected’ arithmetic mean:

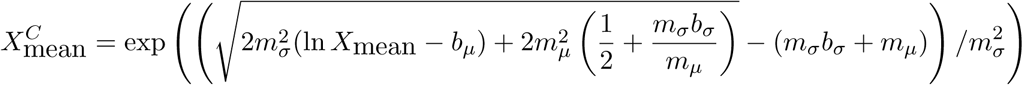

and a ‘corrected’ median:

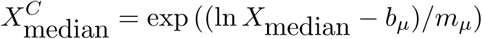

This procedure can be readily adapted for other estimation tasks, distributions of estimates, and estimation biases.

### 2.4. A maximum-likelihood aggregation measure

For this aggregation measure, the full set of estimates is used to form a new collective estimate, rather than just an aggregation measure such as the mean or the median to generate a corrected measure. We again invoke the best-fit relationships in Figure 1c-d, which imply that, for a given actual number of objects *J*, we expect a log-normal distribution described by parameters *µ* = *m_µ_* ln(*J*) + *b_µ_* and *σ* = *m_σ_* ln(*J*) + *b_σ_*. We therefore scan across values of *J* and calculate the likelihood that each associated log-normal distribution generated the given set of estimates. The numerosity that maximizes this likelihood becomes the collective estimate of the true value.

## 3. Results

### 3.1. Quantifying estimation bias

To uncover individual biases in estimation tasks, we first sought to characterize how the distribution of individual estimates changes as a function of the true number of objects *J* (Figure 1a). We performed experiments across a *>*500-fold range of numerosities, from 54 to 27852 objects, with a total of 812 people sampled across the experiments. For all numerosities tested, an approximately log-normal distribution was observed (see Figure 1b for a histogram of an example dataset, Figure S2 for histograms of all other datasets, and Figure S3 for a comparison of the datasets to log-normal distributions). Log-normal distributions can be described by two parameters, *µ* and *σ*, which correspond to the arithmetic mean and standard deviation, respectively, of the normal distribution that results when the original estimates are log-transformed (Figure 1b, inset, and section 1 of the electronic supplementary material on how the maximum-likelihood estimates of *µ* and *σ* were computed for each dataset).

We found that the shape of the log-normal distribution changes in a predictable manner as the numerosity changes. In particular, the two parameters of the log-normal distribution, *µ* and *σ*, both exhibit a linear relationship with the logarithm of the number of objects in the jar (Figure 1c-d). These relationships hold across the entire range of numerosities that we tested (which spans nearly three orders of magnitude). That the parameters of the distribution co-vary closely with numerosity allows us to directly compute how the magnitude of various aggregation measures changes with numerosity, and provides us with information about human estimation behavior which we can exploit to improve the accuracy of the aggregation measures.

### 3.2. Expected error of aggregation measures

We used the maximum-likelihood relationships shown in Figure 1c-d to first compute the expected value of the arithmetic mean, given by exp(*µ* + *σ*^2^*/*2), and the median, given by exp(*µ*), of the log-normal distribution of estimates, across the range of numerosities that we tested empirically (between 54 and 27852 objects). We then compared the magnitude of these two aggregation measures to the true value to identify any systematic biases in these measures (we note that any aggregation measure may be examined in this way, but for clarity here we display just the two most commonly used measures).

Overall, across the range of numerosities tested, we found that the arithmetic mean tended to overestimate, while the median tended to underestimate, the true value (Figure 2a). This is corroborated by our empirical data: for four out of the five datasets, the mean overestimated the true value, while the median underestimated the true value in four of five datasets (Figure 2a). We note that our model predicts qualitatively different patterns for very small numerosities (outside of the range that we tested experimentally). Specifically, in this regime the model predicts that the mean and the median both overestimate the true value, with large relative errors for both measures. However, we expect humans to behave differently when presented with a small number of objects that can be counted directly compared to a large number of objects that could not be easily counted; therefore, we avoid extrapolating our results and apply our model only to the range that we tested experimentally (spanning nearly three orders of magnitude).

**Figure 2:**
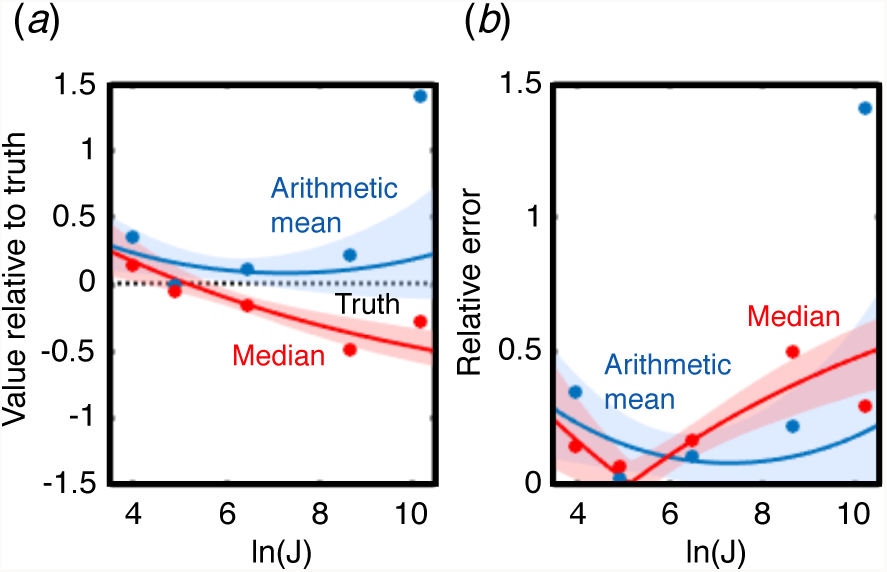
The accuracy of the arithmetic mean and the median. (a) The expected value of the arithmetic mean (blue) and median (red) relative to the true number of objects (black dotted line), as a function of ln(*J*). The relative value is defined as (*X − J*)*/J*, where *X* is the value of the aggregation measure. (b) The relative error of the expected value of the two aggregation measures, defined as *|X − J|/J*. For both panels, solid lines indicate maximum-likelihood values, shaded areas indicate 95% confidence intervals, and solid circles show the empirical values from the five datasets.

That the median tends to underestimate the true value implies that the majority of individuals underestimate the true numerosity. This conforms with the results of other studies demonstrating an underestimation bias in numerosity estimation in humans (*e.g.*, [21–23, 32]). Despite this, the arithmetic mean tends to overestimate the true value because the log-normal distribution has a long tail (Figure 1b), which inflates the mean. Indeed, because the parameter *σ* increases with numerosity, the dispersion of the distribution is expected to increase disproportionally quickly with numerosity, such that the coefficient of variation (the ratio between the standard deviation and the mean of the untransformed estimates) increases with numerosity (Figure S4). This finding differs from other results showing a constant coefficient of variation across numerosities [20, 21]. This contrasting result may be explained by the larger-than-typical range of numerosities that we evaluated here (with respect to previous studies), which improves our ability to detect a trend in the coefficient of variation. Alternatively (and not mutually exclusively), it may result from other studies displaying many numerosities to the same participant, which may cause correlations in a participant’s estimates [21, 22] and reduce variation. By contrast, we only showed a single jar to each participant in our estimation experiments. Overall, the degree of underestimation and overestimation of the median and mean, respectively, was approximately equal across the range of numerosities tested, and we did not detect consistent differences in accuracy between these two aggregation measures (Figure 2b).

### 3.3. Designing and testing aggregation measures that counteract estimation bias

Knowing the expected error of the aggregation measures relative to the true value, we can design new measures to counter this source of collective estimation error. Using this methodology, we specify functional forms of the ‘corrected’ arithmetic mean and the ‘corrected’ median (Methods). In addition to these two adjusted measures, we propose a maximum-likelihood method that uses the full set of estimates, rather than just the mean or median, to locate the numerosity that is most likely to have produced those estimates (Methods). Although applied here to the case of log-normal distributions and particular relationships between numerosity and the parameters of the distributions, our procedure is general and could be used to construct specific corrected measures appropriate for other distributions and relationships, subsequent to empirically characterizing these patterns.

Once the corrected measures have been parameterized for a specific context, they can be applied to a new test dataset to produce an improved collective estimate from that data. However, the three new measures are predicted to have near-zero error only in their expected values, which assumes an infinitely large test dataset (and that the corrected measures have been accurately parameterized). A finite-sized set of estimates, on the other hand, will generally exhibit some deviation from the expected value. It is possible that the measures will produce different noise distributions around the expected value, which will affect their real-world accuracy. To address this, we measured the overall accuracy of the aggregation measures across a wide range of test sample sizes and numerosities, simulating datasets by drawing samples using the maximum-likelihood fits shown in Figure 1c-d. We also conducted a separate analysis, in which we generate test datasets by drawing samples directly from our experimental data, the results of which we include in the electronic supplementary material (see section 2 of the electronic supplementary material for details on both methodologies and for justification of why we chose to include the results from the simulated data in the main text.)

We compared each of the new aggregation measures to the arithmetic mean, the median, and three other ‘standard’ measures that have been described previously in the literature: the geometric mean, the average of the mean and the median, and a trimmed mean (where we remove the smallest 10% of the data, and the largest 10% of the data, before computing the arithmetic mean), in pairwise fashion, calculating the fraction of simulations in which one measure had lower error than the other.

All three new aggregation measures outperformed all of the other measures (Figure 3a, left five columns), displaying lower error in 58–78% of simulations. Comparing the three new measures against each other, the maximum-likelihood measure performed best, followed by the corrected mean, while the corrected median resulted in the lowest overall accuracy (Figure 3a, right three columns). The 95% confidence intervals of the percentages are, at most, *±*1% of the stated percentages (binomial test, *n* = 10000), and therefore the results shown in Figure 3a are all significantly different from chance. The results from our alternate analysis, using samples drawn from our experimental data, are broadly similar, albeit somewhat weaker, than those using simulated data: the corrected median and maximum-likelihood measures still outperformed all of the five standard measures, while the corrected mean outperformed three out of the five standard measures (Figure S5a).

**Figure 3:**
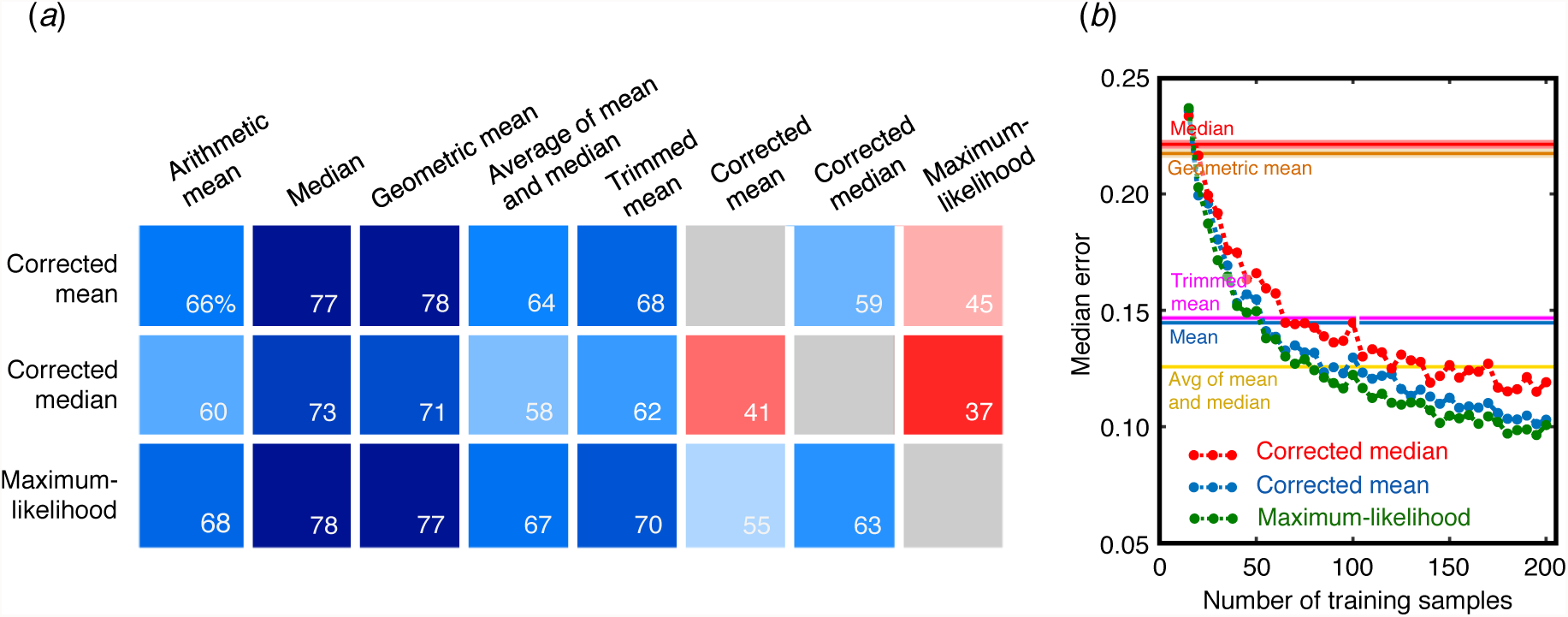
The overall relative performance of the aggregation measures. (a) The percentage of simulations in which the measure indicated in the row was more accurate than the measure indicated in the column. The three new measures are listed in the rows and are compared to all eight measures in the columns. Colors correlate with percentages (blue: *>*50%, red: *<*50%). (b) The median error of the three new aggregation measures (corrected median, dashed red line; corrected mean, dashed blue line; maximum-likelihood measure, dashed green line) as a function of the size of the training dataset. The three new aggregation measures are compared against the arithmetic mean (solid blue), median (solid red), the geometric mean (orange), the average of the mean and the median (yellow), and the trimmed mean (magenta). The 95% confidence interval are displayed for the latter measures, which are not a function of the size of the training dataset.

While the above analysis suggests that the new aggregation measures may be more accurate than many standard measures over a wide range of conditions, it relied on over 800 estimates to parameterize the individual estimation biases. Such an investment to characterize estimation biases may be unfeasible for many applications, so we asked how large of a training dataset is necessary in order to observe improvements in accuracy over the standard measures. To study this, we obtained a given number of estimates from across the range of numerosities, generated a maximum-likelihood regression on that training set, then used that to predict the numerosity of a separate test dataset. As with the previous analysis, we generated the training and test datasets by drawing samples using the maximum-likelihood fits shown in Figure 1c-d, but also conducted a parallel analysis whereby we generated training and test datasets by drawing from our experimental data (section 3 of the electronic supplementary material for details of both methodologies).

We found rapid improvements in accuracy as the size of the training dataset increased (Figure 3b). In our simulations, the maximum-likelihood measure begins to outperform the median and geometric mean when the size of the training dataset is at least 20 samples, the arithmetic mean and trimmed mean after 55 samples, and the average of the mean and median after 80 samples. The corrected mean required at least 105 samples, while the corrected median required at least 175 samples, to outperform the five standard measures. Using samples drawn from our experimental data, our three measures required approximately 200 samples to outperform the five standard measures (Figure S5b). In short, while our method of correcting biases requires parameterizing bias across the entire range of numerosities of interest, our simulations show that much fewer training samples is sufficient for our new aggregation measures to exhibit an accuracy higher than standard aggregation measures.

We next investigated precisely how the size of the test dataset affects accuracy. We defined an ‘error tolerance’ as the maximum acceptable error of an aggregation measure and asked what is the probability that a measure achieves a given tolerance for a particular experiment (the ‘tolerance probability’). As before, we generate test samples by drawing from the maximum-likelihood fits but also perform an analysis drawing from our experimental data (see section 4 of the electronic supplementary material for both methodologies). For all numerosities, the three new aggregation measures tended to outperform the five standard measures if the size of the test dataset is relatively large (Figure 4b-c, Figures S6-S7). However, when the numerosity is large and the size of the test dataset is relatively small, we observed markedly different patterns. In this regime, the relative accuracy of aggregation measures can depend on the error tolerance. For example, for numerosity ln(*J*) = 10, for small error tolerances (*<*0.4), the geometric mean exhibited the lowest tolerance probability across all of the measures under consideration, but for large error tolerances (*>*0.75), it is the most likely to fall within tolerance (Figure 4a). This means that if a researcher wants the collective estimate to be within 40% of the true value (error tolerance of 0.4), then the geometric mean would be the worst choice for small test datasets at large numerosities, but if the tolerance was instead set to 75% of the true value, then the geometric mean would be the best out of all of the measures. These patterns were also broadly reflected in our analysis using samples drawn from our experimental data (Figures S8-S10). Therefore, while the corrected measures should have close to perfect accuracy at the limit of infinite sample size (and perform better than the standard measures overall), there exist particular regimes in which the standard measures may outperform the new measures.

**Figure 4:**
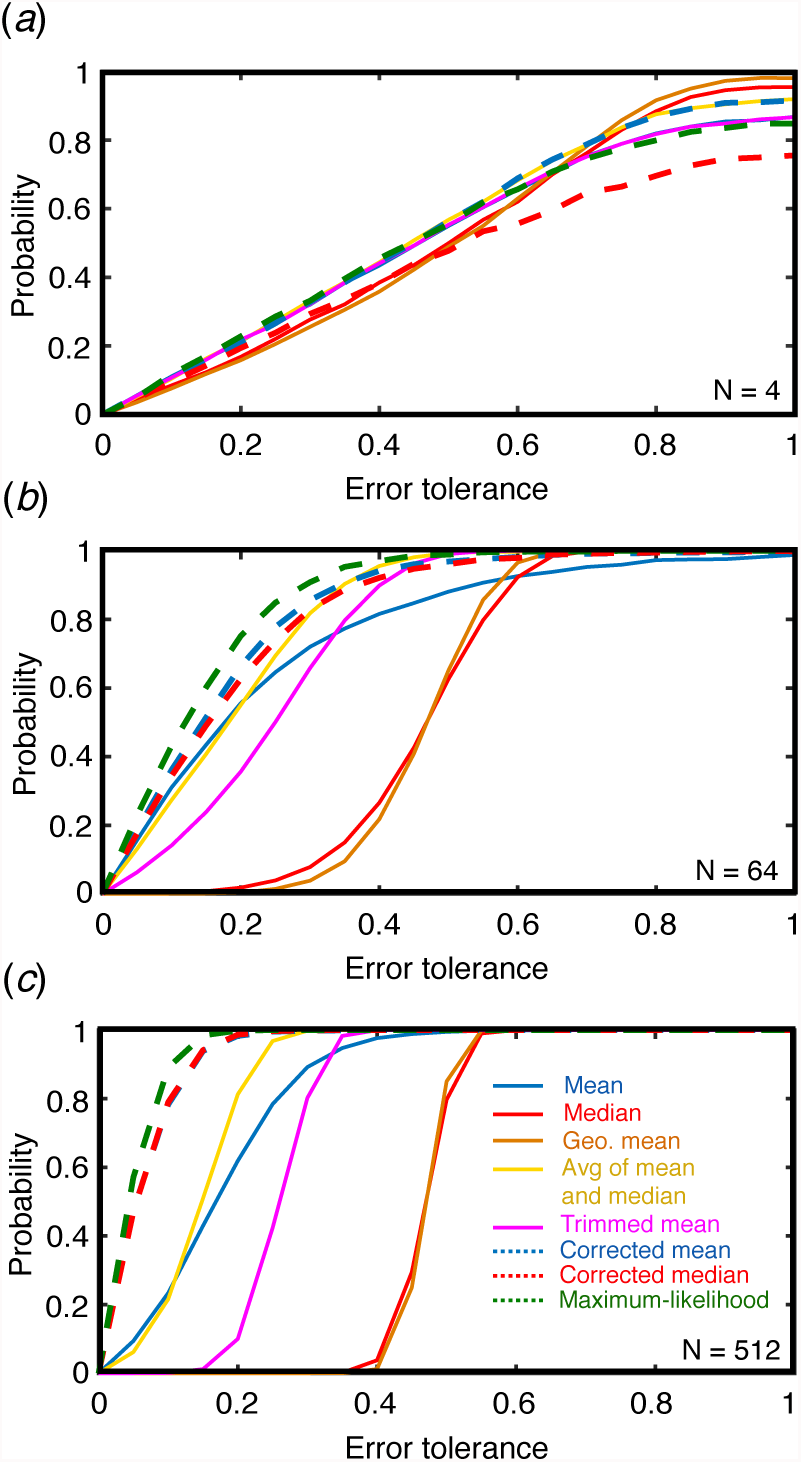
The effect of the test dataset size and error tolerance level on the relative accuracy of the aggregation measures. The probability that an aggregation measure exhibits a relative error (defined as *|X − J|/J*, where *X* is the value of an aggregation measure) less than a given error tolerance, for test dataset size (a) 4, (b) 64, and (c) 512, and numerosity *J* = 22026 (ln(*J*) = 10). In panel (a), the lines for the arithmetic mean and the trimmed mean are nearly identical; in panel (c), the lines for the corrected mean and corrected median are nearly identical.

### 3.4. Quantifying the social influence rules

We then conducted an experiment to quantify the social influence rules that individuals use to update their personal estimate by incorporating information about the estimates of other people (see Methods for details). Briefly, we first allowed participants to make an independent estimate. Then we generated artificial ‘social information’ by selecting a value that was a certain displacement from their first estimate (the ‘social displacement’), and informed the participants that this value was the result of averaging across a certain number of previous estimates (the ‘social group size’). We gave the participants the opportunity to revise their estimate, and we measured how their change in estimate was affected by the social displacement and social group size. By using artificial information and masquerading it as real social information, unlike previous studies, we were able to decouple the effect of social group size, social displacement, and the accuracy of the initial estimate.

We found that a fraction of participants (231 out of 602 participants) completely discounted the social information, meaning that their second estimate was identical to their first. We constructed a two-stage hurdle model to describe the social influence rules by first modeling the probability that a participant utilized or discarded social information, then, for the 371 participants who did utilize social information, we modeled the magnitude of the effect of social information.

A Bayesian approach to fitting a logistic regression model was used to infer whether social displacement (defined as (*S − G*_1_)*/G*_1_, where *S* is the social estimate and *G*_1_ is the participant’s initial estimate), social distance (the absolute value of social displacement), or social group size affected the probability that a participant ignored, or used, social information (see section 5 of the electronic supplementary material for details). Because social distance is a function of social displacement, we did not make inferences about these two variables separately based on their respective credible intervals (coefficient [95% CI]: 0.22 [0.03,0.40] for social displacement and 0.061 [-0.12, 0.24] for social distance). Instead, we graphically interpreted how these two variables jointly affect the probability of changing one’s estimate in response to social information, and overall we found that numerically larger social estimates increased the probability of changing one’s guess, but numerically smaller social estimates decreased that effect (Figure 5a). The probability of using social information did not depend credibly on social group size (−0.045 [−0.18, 0.094]) (Figure 5b). Posterior predictive checks were used to verify the model captured statistical features of the data (Figure S12); see Figure S11a for the posterior distributions.

**Figure 5:**
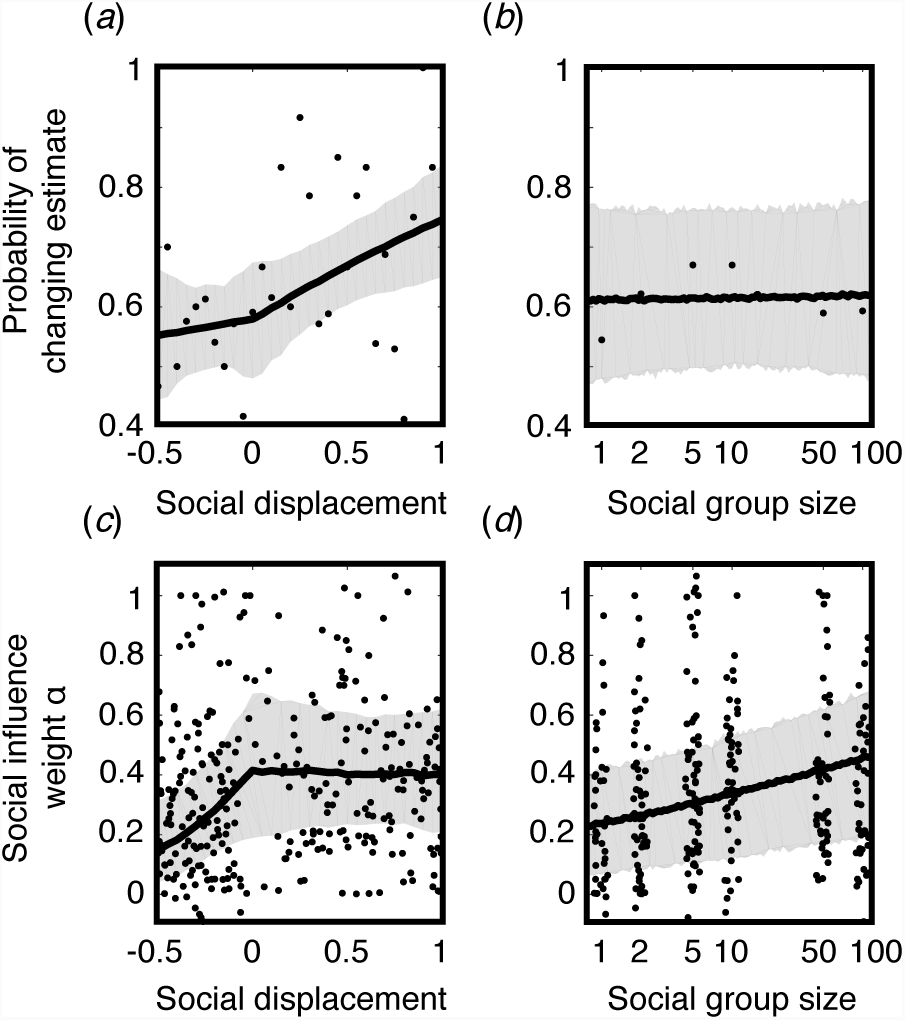
The social influence rules. The probability that an individual is affected by social information as a function of (a) social displacement (the relative displacement of the value of the social information from the participant’s initial estimate) and (b) perceived social group size. The social influence weight *α* for those who used social information as a function of (c) social displacement and (d) social group size. Solid lines: predicted mean value; shaded area: 95% credible interval; circles: the mean of binned data for (a-b) and raw data for (c-d). See Figure S11 for the posterior distributions of each predictor variable. We note that a small fraction of the empirical data extend outside of the bounds of the plots in (c-d); we selected the bounds to more clearly show the patterns of the fitted parameters.

We next modeled the magnitude of the change in estimate, out of the participants who did utilize social information. Following [33], we defined a measure of the strength of social influence, *α*, by considering the logarithm of the participant’s revised estimate, ln(*G*_2_), as a weighted average of the logarithm of the perceived social information, ln(*S*), and the logarithm of the participant’s initial estimate ln(*G*_1_), such that ln(*G*_2_) = *α* ln(*S*) + (1 *− α*) ln(*G*_1_). Here, *α* = 0 indicates that the participant’s two estimates were identical, and therefore the individual was not influenced by social information at all, while *α* = 1 means the participant’s second estimate mirrors the social information. We again used Bayesian techniques to estimate *α* as a normally distributed, logistically transformed linear function of social displacement, social distance, and group size (see section 5 of the electronic supplementary material for details).

Graphically, we found that the social influence weight decreases as the social information is increasingly smaller than the initial estimate but little effect for social information larger than the initial estimate (coeff. [95% CI]: 0.65 [0.28, 1.07] for social displacement and −0.41 [−0.82, −0.0052] for social distance) (Figure 5c). The social influence weight credibly increases with social group size (0.37 [0.17, 0.58]) (Figure 5d). Again, posterior predictive checks revealed that the model generated an overall distribution of social weights consistent with what was found in the data (Figure S13); see Figure 11b for the posterior distributions.

### 3.5. The effect of social influence on the wisdom of crowds

If individuals share information with each other before their opinions are aggregated, then the independent, log-normal distribution of estimates will be altered. Since individuals take a form of weighted average of their own estimate and perceived social information, the distribution of estimates should converge towards intermediate values. However, it is not clear what effect the observed social influence rules have on the value, or accuracy, of the aggregation measures [34]. In particular, since the new aggregation measures introduced here were parameterized on independent estimates unaltered by social influence, their performance may degrade when individuals share information with each other.

We simulated several rounds of influence using the rules that we uncovered, using a fully connected social network (each individual was connected all other individuals), in order to identify measures that may be relatively robust to social influence (see section 6 of the electronic supplementary material). We used two alternate assumptions about how a set of estimates is averaged, either by the individual or by an external agent, before being presented as social information (the ‘individual aggregation measure’), using either the geometric mean or the arithmetic mean (see section 7 of the electronic supplementary material). While the maximum-likelihood measure generally performed the best in the absence of social influence (Figure 3), this measure was highly susceptible to the effects of social influence, particularly at large numerosities (Figure 6). By contrast, the corrected mean was remarkably robust to social influence, across numerosities, and for both individual aggregation measures, while exhibiting nearly the same accuracy as the maximum-likelihood measure in the absence of social influence.

**Figure 6:**
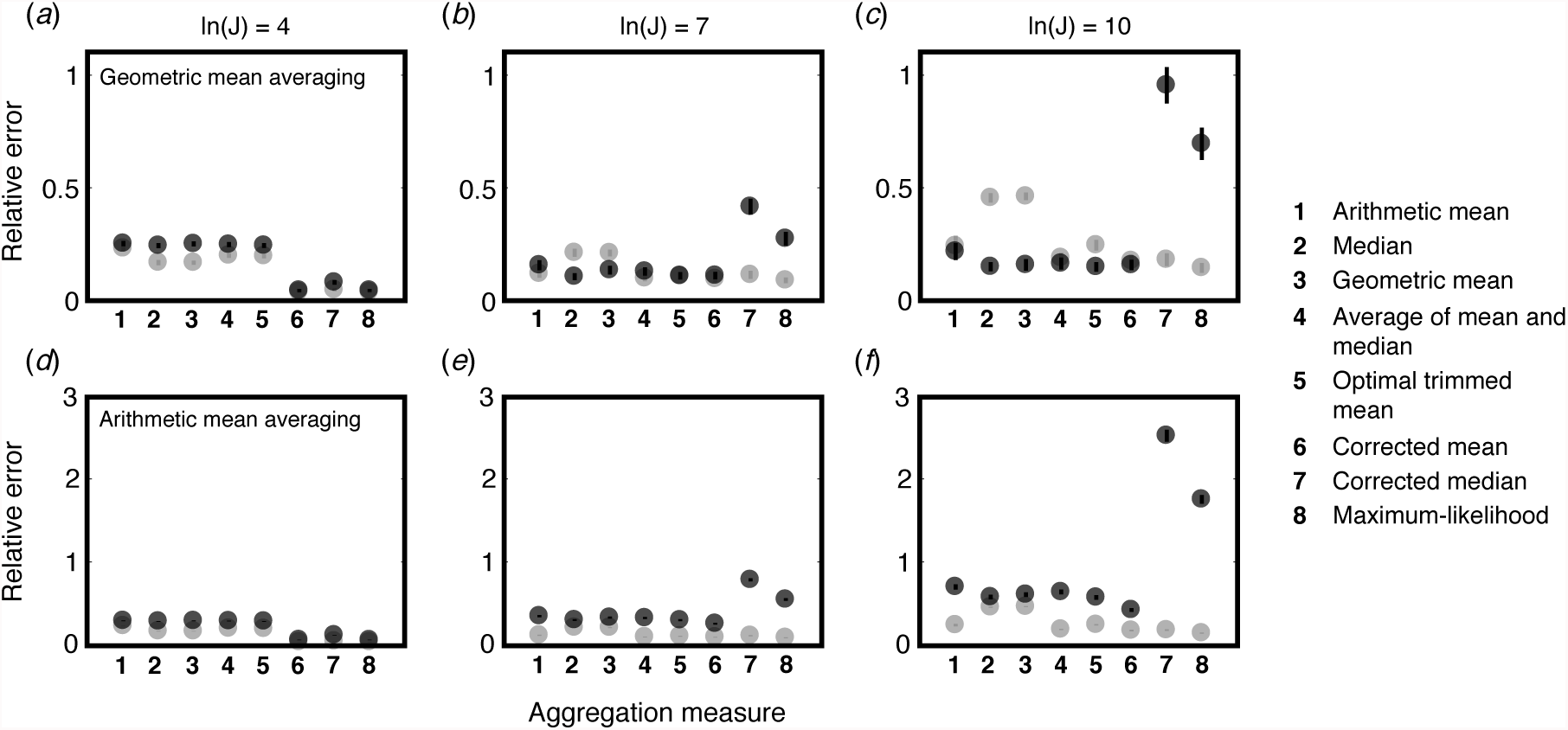
The robustness of aggregation measures under social influence. The relative error of the eight aggregation measures without social influence (light gray circles) and after ten rounds of social influence (dark gray circles) when (a-c) individuals internally take the geometric mean of the social information that they observe, or when (d-f) individuals internally take the arithmetic mean of the social information, for numerosity ln(*J*) = 4 (a,d), ln(*J*) = 7 (b,e), and ln(*J*) = 10 (c,f). Circles show the mean relative error across 1000 replicates, error bars show twice the standard error. The error bars are often smaller than the size of the corresponding circles, and where some gray circles are not visible, they are nearly identical to the corresponding black circles.

## 4. Discussion

While the wisdom of crowds has been documented in many human and non-human contexts, the limits of its accuracy are still not well understood. Here we demonstrated how, why, and when collective wisdom may break down by characterizing two major sources of error, individual (estimation bias) and social (information sharing). We revealed the limitations of some of the most common averaging measures and introduced three novel measures that leverage our understanding of these sources of error to improve the wisdom of crowds.

In addition to the conclusions and recommendations drawn for numerosity estimation, the methods described here could be applied to a wide range of estimation tasks. Estimation biases and social influence are ubiquitous, and estimation tasks may cluster into broad classes that are prone to similar biases or social rules [35]. For example, the distribution of estimates for many tasks are likely to be log-normal in nature [36], while others may tend to be normally distributed. Indeed, there is evidence that counteracting estimation biases can be a successful strategy to improve estimates of probabilities [37–39], city populations [40], movie box office returns [40], and engineering failure rates [41].

Furthermore, the social influence rules that we identified empirically are similar to general models of social influence, with the exception of the effect of the social displacement that we uncovered. This asymmetric effect suggests that a focal individual was more strongly affected by social information that was larger in value relative to the focal individual’s estimate compared to social information that was smaller than the individual’s estimate. The observed increase in the coefficient of variation as numerosity increased (Figure S4b) may suggest that one’s confidence about one’s own estimate decreases as numerosity increases, which could lead to an asymmetric effect of social displacement. Other estimation contexts in which confidence scales with estimation magnitude could yield a similar effect. This effect was combined with a weaker negative effect of the social distance, which is reminiscent of ‘bounded confidence’ opinion dynamics models (*e.g.*, [42–44]), whereby individuals weigh more strongly social information that is similar to their own opinion. By carefully characterizing both the *individual* estimation biases and *collective* biases generated by social information sharing, our approach allows us to counteract such biases, potentially yielding significant improvements when aggregating opinions across other domains.

Other approaches have been used to improve the accuracy of crowds. One strategy is to search for ‘hidden experts’ and weigh these opinions more strongly [3, 33, 45–48]. While this can be effective in certain contexts, we did not find evidence of hidden experts in our data. Comparing the group of individuals who ignored social information and those who utilized social information, the two distribution of estimations were not significantly different (*P* = 0.938, Welch’s t-test on the log-transformed estimates), and the arithmetic mean, the median, nor our three new aggregation measures were significantly more accurate across the two groups (Figure S14). Furthermore, searching for hidden experts requires additional information about the individuals (such as propensity to use social information, past performance, or confidence level). Our method does not require any additional information about each individual, only knowledge about statistical tendencies of the population at large (and relatively few samples may be needed to sufficiently parameterize these tendencies).

Further refinement of our methods is possible. In cases where the underlying social network is known [49, 50], or where individuals vary in power or influence [51], simulation of social influence rules on these networks could lead to a more nuanced understanding of the mapping between individual to collective estimates. In addition, aggregation measures can be generalized in a straightforward manner to calculate confidence intervals, in which an estimate range is generated that includes the true value with some probability. To improve the accuracy of confidence intervals, information about the sample size and other features that we showed to be important can be included.

In summary, counteracting estimation biases and social influence may be a simple, general, and computationally efficient strategy to improve the wisdom of crowds.

## 5. Competing interests

We have no competing interests.

## 6. Author Contributions

ABK, AB, and IDC designed the experiments. ABK, AB, ATH, and MJL performed the experiments. ABK, AB, JB-C, CCI, and XG analyzed the data. ABK, AB, and IDC wrote the paper.

## 7. Acknowledgements

We thank Stefan Krause, Jens Krause, Andrew King, and Michael J. Mauboussin for contributing datasets to this study, and Mirta Galesic for providing feedback on the manuscript.

## 8. Data Accessibility

Datasets are available in the electronic supplementary material.

## 9. Ethics

The experimental procedures were approved by the Princeton University and Santa Fe Institute ethics committees.

## 10. Funding Statement

ABK was supported by a James S. McDonnell Foundation Postdoctoral Fellowship Award in Studying Complex Systems. AB was supported by an SFI Omidyar Postdoctoral Fellowship and a grant from the Templeton Foundation. CCI was supported by a NERC Independent Research Fellowship NE/K009370/1. IDC acknowledges support from NSF (PHY-0848755, IOS-1355061, EAGER-IOS-1251585), ONR (N00014-09-1-1074, N00014-14-1-0635), ARO (W911NG-11-1-0385, W911NF-14-1-0431), and the Human Frontier Science Program (RGP0065/2012).

## References

[1] J. Surowiecki, The Wisdom of the Crowds: Why the Many are Smarter than the Few, Little Brown, 2004.

[2] F. Galton, Vox populi, Nature 75 (1907) 450–451.

[3] D. Prelec, H. Seung, J. McCoy, A solution to the single-question crowd wisdom problem, Nature 541 (2017) 532–535.

[4] B. Bahrami, K. Olsen, P. Latham, A. Roepstorff, G. Rees, C. Frith, Optimally interacting minds, Science 329 (2010) 1081–1085.

[5] S. Krause, R. James, J. Faria, G. Ruxton, J. Krause, Swarm intelligence in humans: diversity can trump ability, Animal Behaviour 81 (2011) 941–948.

[6] A. King, L. Cheng, S. Starke, J. Myatt, Is the true ‘wisdom of the crowd’ to copy successful individuals?, Biology Letters 8 (2011) 197–200.

[7] D. Sumpter, J. Krause, R. James, I. Couzin, A. Ward, Consensus decision making by fish, Current Biology 18 (2008) 1773–1777.

[8] A. Ward, J. Herbert-Read, D. Sumpter, J. Krause, Fast and accurate decisions through collective vigilance in fish shoals, Proc Natl Acad Sci USA 108 (2011) 2312–2315.

[9] A. Ward, D. Sumpter, I. Couzin, P. Hart, J. Krause, Quorum decision-making facilitates information transfer in fish shoals, Proc Natl Acad Sci USA 105 (2008) 6948–6953.

[10] C. C. Ioannou, Swarm intelligence in fish? the difficulty in demonstrating distributed and self-organised collective intelligence in (some) animal groups, Behavioural processes 141 (2017) 141–151.

[11] T. Sasaki, B. Granovskiy, R. Mann, D. Sumpter, S. Pratt, Ant colonies outperform individuals when a sensory discrimination task is difficult but not when it is easy, Proc Natl Acad Sci USA 110 (2013) 13769–13773.

[12] T. Sasaki, S. Pratt, Emergence of group rationality from irrational individuals, Behav Ecol 22 (2011) 276–281.

[13] S. Tamm, Bird orientation: single homing pigeons compared to small flocks, Behav Ecol Sociobiol 7 (1980) 319–322.

[14] A. Simons, Many wrongs: the advantage of group navigation, Trends Ecol Evol 19 (2004) 453–455.

[15] N. Condorcet, Essai sur l’application de l’analyse à la probabilité des décisions rendues à la pluralité de voix, Paris, 1785.

[16] D. Kahneman, Thinking, Fast and Slow, Straus and Giroux, 2011.

[17] R. Nickerson, Confirmation bias: a ubiquitous phenomenon in many guises, Review of General Psychology 2 (1998) 175–220.

[18] M. Haselton, D. Nettle, The paranoid optimist: an integrative evolutionary model of cognitive biases, Personality and Social Psychology Review 10 (2006) 47–66.

[19] S. Stevens, On the psychophysical law, The Psychological Review 64 (1957) 153–181.

[20] J. Whalen, C. Gallistel, R. Gelman, Nonverbal counting in humans: the psychophysics of number representation, Psychological Science 10 (1999) 130–137.

[21] V. Izard, S. Dehaene, Calibrating the mental number line, Cognition 106 (2008) 1221–1247.

[22] L. E. Krueger, Single judgments of numerosity, Attention, Perception, & Psychophysics 31 (1982) 175–182.

[23] L. E. Krueger, Perceived numerosity: A comparison of magnitude production, magni-tude estimation, and discrimination judgments, Attention, Perception, & Psychophysics 35 (1984) 536–542.

[24] L. Krueger, Reconciling fechner and stevens: toward a unified psychophysical law, Behavioral and Brain Sciences 12 (1989) 251–320.

[25] F. Galton, One vote, one value, Nature 75 (1907) 414.

[26] J. Lorenz, H. Rauhut, F. Schweitzer, D. Helbing, How social influence can undermine the wisdom of crowd effect, Proc Natl Acad Sci USA 108 (2011) 9020–9025.

[27] M. Lobo, D. Yao, Human judgement is heavy tailed: Empirical evidence and implications for the aggregation of estimates and forecasts, Fontainebleau: INSEAD, 2010.

[28] J. Armstrong, Combining forecasts. Armstrong, JS, ed. Principles of Forecasting: A Handbook for Researchers and Practitioners, Kluwer, New York, 2001.

[29] A. Kao, N. Miller, C. Torney, A. Hartnett, I. Couzin, Collective learning and optimal consensus decisions in social animal groups, PLoS Computational Biology 10 (2014) e1003762.

[30] C. Wagner, C. Schneider, S. Zhao, H. Chen, The wisdom of reluctant crowds, Proceedings of the 43rd Hawaii International Conference on System Sciences, 2010.

[31] M. Mauboussin, Explaining the wisdom of crowds, Legg Mason Capital Management White Paper, 2007.

[32] S. Kemp, Estimating the sizes of sports crowds, Perceptual and motor skills 59 (1984) 723–729.

[33] G. Madirolas, G. de Polavieja, Improving collective estimations using resistance to social influence, PLoS Comput Biol 11 (2015) e1004594.

[34] B. Golub, M. Jackson, Naïve learning in social networks and the wisdom of crowds, American Economic Journal: Microeconomics 2 (2010) 112–149.

[35] M. Steyvers, B. Miller, Cognition and collective intelligence, Handbook of Collective Intelligence (2015) 119.

[36] S. Dehaene, V. Izard, E. Spelke, P. Pica, Log or linear? distinct intuitions of the number scale in western and amazonian indigene cultures, Science 320 (2008) 1217–1220.

[37] B. M. Turner, M. Steyvers, E. C. Merkle, D. V. Budescu, T. S. Wallsten, Forecast aggregation via recalibration, Machine learning 95 (2014) 261–289.

[38] M. D. Lee, I. Danileiko, Using cognitive models to combine probability estimates, Judgment and Decision Making 9 (2014) 259.

[39] V. A. Satopää, J. Baron, D. P. Foster, B. A. Mellers, P. E. Tetlock, L. H. Ungar, Combining multiple probability predictions using a simple logit model, International Journal of Forecasting 30 (2014) 344–356.

[40] A. Whalen, S. Yeung, Using ground truths to improve wisdom of the crowd estimates, Proceedings of the Annual Cognitive Science Society Meeting (2015).

[41] E. Merkle, M. Steyvers, A psychological model for aggregating judgments of magnitude, Social computing, behavioral-cultural modeling and prediction (2011) 236–243.

[42] R. Hegselmann, U. Krause, Opinion dynamics and bounded confidence models, analysis, and simulation, J Artif Soc Soc Simulat 5 (2002).

[43] G. Deffuant, D. Neau, F. Amblard, G. Weisbuch, Mixing beliefs among interacting agents, Advances in Complex Systems 3 (2000) 87–98.

[44] G. Deffuant, F. Amblard, G. Weisbuch, T. Faure, How can extremism prevail? a study based on the relative agreement interaction model, Journal of artificial societies and social simulation 5 (2002).

[45] A. Koriat, When are two heads better than one and why?, Science 336 (2012) 360–362.

[46] S. Hill, N. Ready-Campbell, Expert stock picker: the wisdom of (experts in) crowds, International Journal of Electronic Commerce 15 (2011) 73–102.

[47] J. Whitehill, T. Wu, J. Bergsma, J. Movellan, P. Ruvolo, Whose vote should count more: optimal integration of labels from labelers of unknown expertise, Advances in Neural Information Processing Systems 22 (2009) 2035–2043.

[48] D. V. Budescu, E. Chen, Identifying expertise to extract the wisdom of crowds, Management Science 61 (2014) 267–280.

[49] M. Jönsson, U. Hahn, E. Olsson, The kind of group you want to belong to: Effects of group structure on group accuracy, Cognition 142 (2015) 191–204.

[50] M. Moussaïd, S. Herzog, J. Kämmer, R. Hertwig, Reach and speed of judgment propagation in the laboratory, Proc Natl Acad Sci USA 114 (2017) 4117–4122.

[51] J. Becker, D. Brackbill, D. Centola, Network dynamics of social influence in the wisdom of crowds, Proc Natl Acad Sci USA 114 (2017) E5070–E5076.

