## Supplementary Materials for "Counteracting estimation bias and social influence to improve the wisdom of crowds"

Electronic supplementary material for "Counteracting estimation bias and social influence to improve the wisdom of crowds" by Kao AB, Berdahl AM, Hartnett AT, Lutz MJ, Bak-Coleman JB, Ioannou CC, Giam X, and Couzin ID in *Journal of the Royal Society Interface*

### SI.1. Calculating maximum-likelihood fits from the datasets

For our five original datasets and six full datasets from the literature (where the individual estimates were available), the maximum-likelihood estimates and 95% confidence intervals for the parameters $\mu$ and $\sigma$ were computed directly. For the two datasets from the literature for which only summary statistics of the distribution were available, we estimated the parameters $\mu$ and $\sigma$ from the arithmetic mean $M$ and standard deviation $S$ of the data using the equations

$$\sigma = \sqrt{\ln(\exp(\ln S^2 - 2\ln M) + 1)}$$

and

$$\mu = \ln M - \frac{1}{2}\sigma^2$$

To generate confidence intervals for these parameter estimates, we used parametric bootstrapping [1] by creating 10000 sets of estimates of the same size as the original datasets using the parameter estimates $\mu$ and $\sigma$, calculating the $M'$ and $S'$ from each set, and re-calculating the estimates $\mu'$ and $\sigma'$ from $M'$ and $S'$. We then calculated the 95% confidence interval as the range that captures 95% of the estimates of $\mu'$ and $\sigma'$.

### SI.2. Comparing the overall accuracy of the aggregation measures

*Using simulated data*

In this analysis, we generated 10000 sets of estimates. For each test dataset $s$, we selected a sample size $p_s$, where $\log_2(p_s)$ was randomly drawn from a uniform distribution on the interval $[2, 9]$, and a numerosity $j_s$, where $\ln(j_s)$ was randomly drawn from a uniform distribution on the interval $[4, 10]$. $p_s$ estimates were then drawn from the log-normal distribution associated with numerosity $j_s$, based on the maximum-likelihood lines shown in Figure 1c-d in the main text (we assume that this is the 'true' underlying human behavior). We calculated each of the aggregation measures using these $p_s$ estimates. For this analysis, we assumed that our training dataset was the complete set of estimates that we collected across all of our experiments, such that the maximum-likelihood lines shown in Figure 1c-d in the main text were used to compute the 'corrected' aggregation measures. We compared each of the new aggregation measures to all of the other measures in pairwise fashion, respectively for each of the 10000 sets of estimates, and calculated the fraction of simulations for which one measure had lower error than the other.

*Using empirical data*

Here we also generated 10000 sets of estimates. To generate each test dataset, we randomly selected one of our five experimental datasets, and from that experimental dataset, randomly selected 10 estimates without replacement. Our training dataset comprised the remaining estimates, across all of our experimental datasets, that were not included in the test dataset. A new maximum-likelihood fit was computed for this training dataset to estimate the 'true' behavior, and the results from this new fit were used to compute the corrected aggregation measures. All of the aggregation measures were then computed for the 10 estimates comprising the test dataset. As with the simulated data, we then compared each of the new aggregation measures to all of the other measures in pairwise fashion, respectively for each of the 10000 sets of estimates, and calculated the fraction of simulations for which one measure had lower error than the other.

*Justification for displaying the results from the simulated data in the main text*

For our analysis using empirical data, we restricted the size of our test datasets to 10 estimates because our smallest experimental dataset, corresponding to the smallest numerosity, comprised 36 estimates. If we allocated a large fraction of that dataset to the test set, then the training set would be comprised almost entirely of estimates from larger numerosities, and our prediction of the test set would require extrapolating outside of the range of the training set. To avoid this problem, we limited the size of the test dataset to be much smaller than the size of the smallest experimental dataset; however, this limited our ability to explore the behavior of the aggregation measures across a larger range of sizes of test datasets. Because of our ability to explore a much larger range of sizes of test datasets with simulated data, we chose to display the results of the simulated data in the main text, and the results of the empirical data here in the electronic supplementary material.

### SI.3. Testing how the accuracy of the new aggregation measures improves with the size of the training dataset

*Using simulated data*

In this analysis, we varied the size of the training dataset from 15 to 200 samples. For a given training set size $r$, we selected $r$ numerosities, where each numerosity $\ln(j_i)$ was randomly drawn from a uniform distribution on the interval $[4, 10]$. We generated one estimate for each numerosity, drawn from the log-normal distribution associated with that numerosity, based on the maximum-likelihood lines shown in Figure 1c-d in the main text. We then performed a maximum-likelihood regression using these $r$ estimates in order to estimate the underlying behavior (*i.e.*, reproducing Figure 1c-d but using just the $r$ training samples, in order to generate corrected aggregation measures for the test dataset later). We next created one test dataset by selecting a sample size $p_T$, where $\log_2(p_T)$ was randomly drawn from a uniform distribution on the interval

$[2, 9]$, and a numerosity $J_T$, where $\ln(J_T)$ was randomly drawn from a uniform distribution on the interval $[4, 10]$. $p_T$ estimates were then drawn from the the log-normal distribution associated with numerosity $J_T$, based on the maximum-likelihood lines shown in Figure 1c-d in the main text. We then predicted the numerosity of the test set $J_T$ using the aggregation measures and the maximum-likelihood parameters derived from the training round. This process was repeated 2000 times for each training set size.

*Using empirical data*

Here we also varied the size of the training dataset from 15 to 200 samples. To generate each test dataset, we randomly selected one of our five experimental datasets, and from that experimental dataset, randomly selected 10 estimates without replacement. To generate a corresponding training dataset, for a given training set size $r$, we selected $r$ estimates out of the remaining estimates, across all of the experimental datasets, that were not included in the test dataset (without replacement). A new maximum-likelihood fit was computed for this training dataset to estimate the 'true' behavior, and the results from this new fit were used to compute the corrected aggregation measures. All of the aggregation measures were then computed for the 10 estimates comprising the test dataset. This process was repeated 2000 times for each training set size.

### SI.4. Testing how the accuracy of the aggregation measures changes with the size of the test dataset

*Using simulated data*

In this analysis, we examined three test dataset sizes $p$, where $\log_2(p) = \{2, 6, 9\}$ estimates, and three numerosities $J$, where $\ln(J) = \{4, 7, 10\}$. For each combination of test dataset size $p$ and numerosity $J$, we generated 10000 sets of estimates by drawing $p$ estimates from the log-normal distribution associated with numerosity $J$, based on the maximum-likelihood lines shown in Figure 1c-d in the main text (we assume that this is the 'true' underlying human behavior). We then calculated the relative error, given by $|X - J|/J$ (where $X$ is the value of an aggregation measure) for each of the aggregation measures for each set of estimates and computed the fraction of the 10000 sets of estimates that exhibited a relative error smaller than a given error tolerance. For this analysis, we assumed that our training dataset was the complete set of estimates that we collected across all of our experiments, such that the maximum-likelihood lines shown in Figure 1c-d in the main text were used to compute the 'corrected' aggregation measures.

*Using empirical data*

Here we also examined three test dataset sizes $p$, where $\log_2(p) = \{2, 6, 9\}$ estimates, for three of our experimental datasets, corresponding to numerosities $J = \{54, 659, 27852\}$ ($\ln J = \{3.99, 6.49, 10.23\}$). For each combination of test dataset size $p$ and numerosity $J$ we generated

10000 sets of estimates by drawing $p$ estimates, with replacement, from the experimental dataset corresponding to that numerosity. We then calculated the relative error, given by $|X - J|/J$ (where $X$ is the value of an aggregation measure) for each of the aggregation measures for each set of estimates and computed the fraction of the 10000 sets of estimates that exhibited a relative error smaller than a given error tolerance. We assumed that our training dataset was the complete set of estimates that we collected across all of our experiments, such that the maximum-likelihood lines shown in Figure 1c-d in the main text were used to compute the 'corrected' aggregation measures.

#### SI.5. Modeling the social influence rules

In our two-stage hurdle Bayesian model [2], we first modeled the probability an individual discarded or utilized social information. Then, out of those who utilized social information, we modeled the magnitude of the effect of social information.

We assumed that whether an individual discarded or utilized social information is a Bernoulli random variable with probability $p$ of utilizing social information. We described the probability $p$ with a logistic regression model, such that $p = \text{logit}^{-1}(\beta_{1,0} + \beta_{1,D}D + \beta_{1,|D|}|D| + \beta_{1,N}\ln N)$, where $D$ is the 'social displacement' $D = (S - G_1)/G_1$ ($S$ is the value of the social information that the participant perceives and $G_1$ is the participant's initial estimate), $|D|$ is the 'social distance'; $N$ is the group size perceived by the participant (these variables were standardized before running the model); $\beta_{1,D}$, $\beta_{1,|D|}$, and $\beta_{1,N}$ are the respective coefficients for the explanatory variables; and $\beta_{1,0}$ is the intercept (the intercept sets a baseline propensity to utilize social information in the absence of other cues). We used generic uninformative priors (Gaussian distributions with mean 0 and variance 1000) for all coefficients. The model coefficients were fit using JAGS [3] via PyJAGS version 1.2.2 using Python version 2.7.14, running the model until convergence was achieved. The model was then used to simulate data, and the probability of switching as a function of social distance was compared between the data and the model (Figure S12).

Then, for those participants who changed their estimate, we modeled a participant's final estimate $G_2$ as a normally distributed variable with a mean of $\exp(\alpha \ln(S) + (1 - \alpha)\ln(G_1))$ and standard deviation $\sigma_{G_2}$, where, as before, $S$ is the value of the social information that the participant perceives and $G_1$ is the participant's initial estimate. The social influence weight $\alpha$ was considered to be normally distributed about a mean $\mu_\alpha$ and standard deviation $\sigma_\alpha$. To constrain the mean $\mu_\alpha$ between 0 and 1, we modeled it as a logistic function, such that $\mu_\alpha = \text{logit}^{-1}(\beta_{2,0} + \beta_{2,D}D + \beta_{2,|D|}|D| + \beta_{2,N}\ln N)$. We used generic uninformative priors (Gaussian distributions with mean 0 and variance 1000) for all coefficients, and gamma prior distributions ($a = 0.1, b = 0.1$) for the standard deviations $\sigma_{G_2}$ and $\sigma_\alpha$. Model fit was evaluated by examining the distributions of observed and estimated $\alpha$ values (Figure S13).

#### SI.6. Simulating the effect of social influence on the wisdom of crowds

We simulated multiple rounds of social influence for a group of 64 individuals estimating the number of objects in a jar for three numerosities, $\ln(J) = 4$ (55), $\ln(J) = 7$ (1097), and $\ln(J) = 10$ (22026). The simulated individuals began with independent estimates of the numerosity by drawing estimates from the log-normal distributions described by the best-fit lines of Figure 1c-d in the main text for that numerosity. We constructed a fully connected social network where each individual was connected to all other individuals in the group. At each round in the simulation, each individual updated its estimate following our empirically-parameterized social influence model (Figure 5 in the main text and Figure S11). The social information that the selected individual used to update its estimate was either the geometric mean or the arithmetic mean of the estimates of the rest of the group in the previous round (see section 7 below). This process was repeated for ten rounds of social influence. We then calculated the aggregation measures at the start and at the end of the simulation and compared them to the true value. We repeated the simulation 1000 times for each numerosity and for each individual aggregation measure.

#### SI.7. The individual aggregation measure

In our social influence model, we assume that the social information is a single number that represents the average of one or more estimates. In some real world cases, an external mechanism may average the estimates, so that an individual is presented with a single number (such as for online recommendation services). In other cases, the individual may be presented with the full list of estimates, and she must mentally average across the list. It is not well understood how individuals mentally average a set of estimates presented as social information. Theoretical work suggests that humans and other animals may take the geometric mean of a set of estimates [4]. However, in another study [5], researchers presented subjects with either the arithmetic mean of the group's estimates or the full set of estimates and observed how the distribution of the subjects' estimates was altered. The results across these two conditions were highly similar, which may suggest that humans approximately take the mean of a set of estimates. Consequently, we considered both the geometric mean and the arithmetic mean as possible individual aggregation measures.

### References

[1] A. Davison, D. Hinkley, Bootstrap Methods and their Application, Cambridge University Press, 1997.

[2] X. Giam, L. Mani, L. P. Koh, H. T. Tan, Saving tropical forests by knowing what we consume, Conservation Letters 9 (2016) 267–274.

[3] K. Hornik, F. Leisch, A. Zeileis, Jags: A program for analysis of bayesian graphical models using gibbs sampling, in: Proceedings of DSC, volume 2, pp. 1–1.

[4] G. Madirolas, G. de Polavieja, Improving collective estimations using resistance to social influence, PLoS Comput Biol 11 (2015) e1004594.

[5] J. Lorenz, H. Rauhut, F. Schweitzer, D. Helbing, How social influence can undermine the wisdom of crowd effect, Proc Natl Acad Sci USA 108 (2011) 9020–9025.

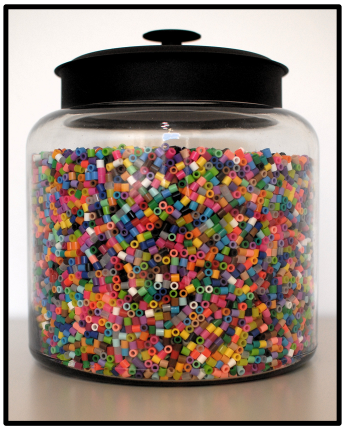

J = 27852 objects
ln(J) = 10.2

Figure S1: **An example jar containing 27852 objects.** The same beads were also used for the jar containing 5897 objects, with a similarly shaped jar that was scaled so that it remained approximately 90% full. For the other three numerosities, we used gumballs such as those shown in Figure 1a in the main text, where for each numerosity, a different jar was used such that each jar was approximately 90% full.

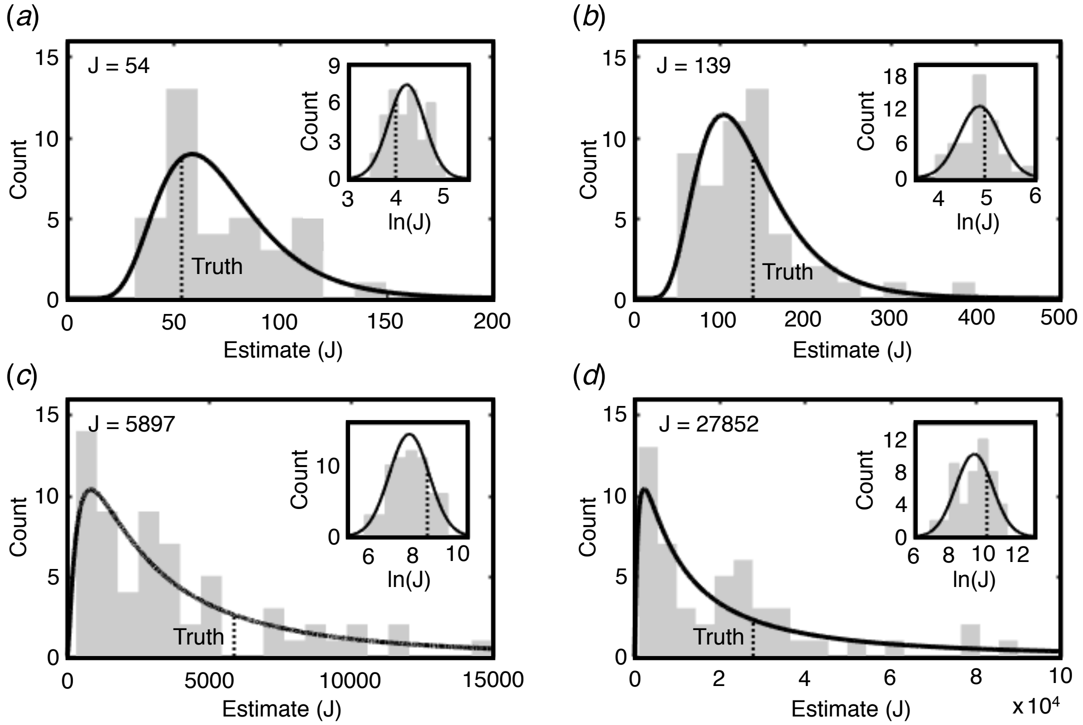


Figure S2: **The distribution of estimates in the datasets collected in this study.** In addition to the distribution of estimates collected using a jar containing 659 objects, shown in Figure 1b in the main text, we collected four more sets of estimates, using jars containing (a) 54, (b) 139, (c) 5897, and (d) 27852 objects. The histogram of estimates is shown in gray, the maximum-likelihood log-normal distribution in black, and the true number of objects shown as a vertical dashed line. Insets show the log-transformed estimates along with the maximum-likelihood normal distribution and the true value.

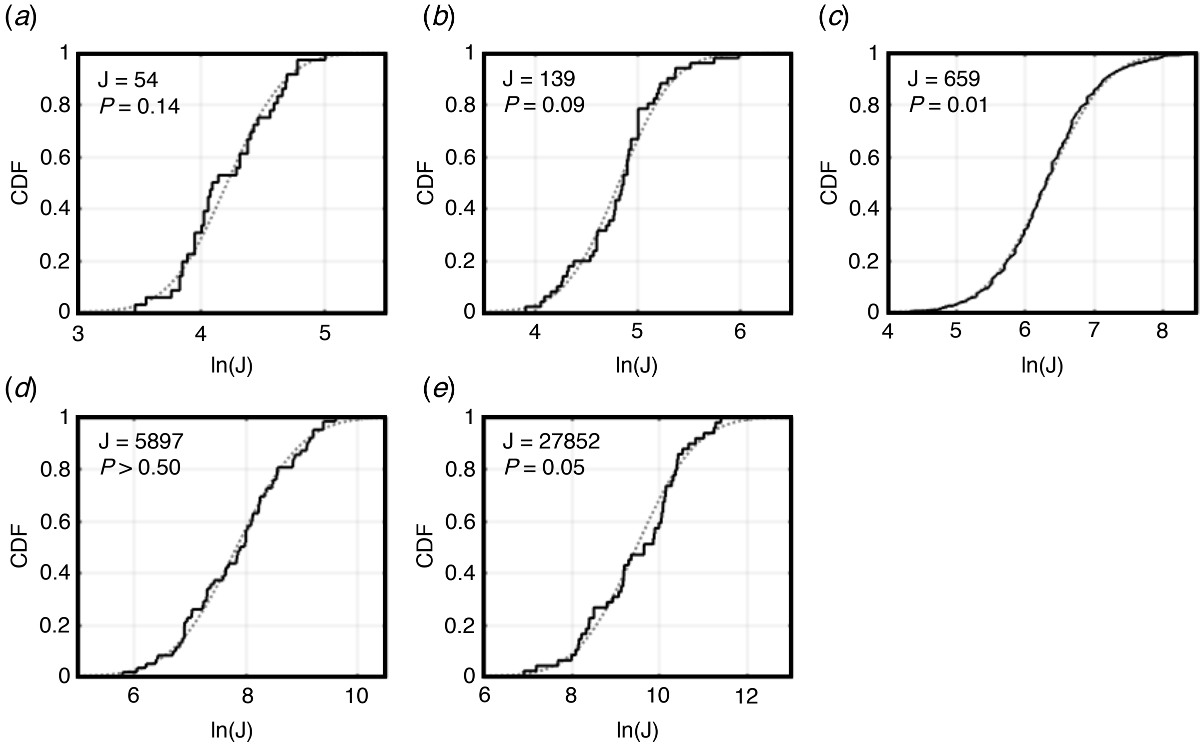


Figure S3: **Statistical comparison between the empirical distributions of estimates and log-normal distributions.** Shown are the empirical cumulative distribution functions (CDF) of the log-transformed estimates (black lines) collected using jars containing (a) 54, (b) 139, (c) 659, (d) 5897, and (e) 27852 objects, with the CDF of the normal distribution, with mean and standard deviation equal to that of the respective set of estimates, shown as dashed gray lines. P-values are from a Kolmogorov-Smirnov test for normality with Lilliefors correction.

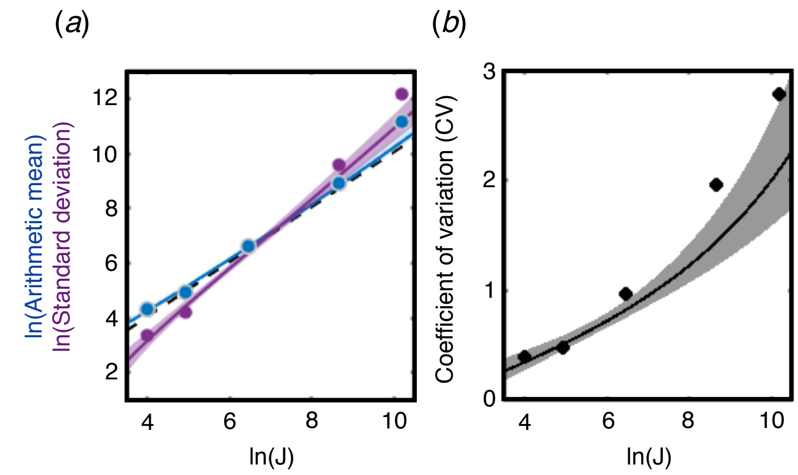


Figure S4: **The arithmetic mean, standard deviation, and coefficient of variation all increase with numerosity.** (a) The logarithm of the arithmetic mean (blue) and the logarithm of the standard deviation (purple) of the estimates, as a function of $\ln(J)$. Black dashed line shows the $y = x$ line. (b) The expected value of the coefficient of variation, defined as the ratio of the standard deviation to the arithmetic mean. For both panels, solid lines indicate maximum-likelihood values, shaded areas indicate 95% confidence intervals, and solid circles show the empirical values from the five datasets.

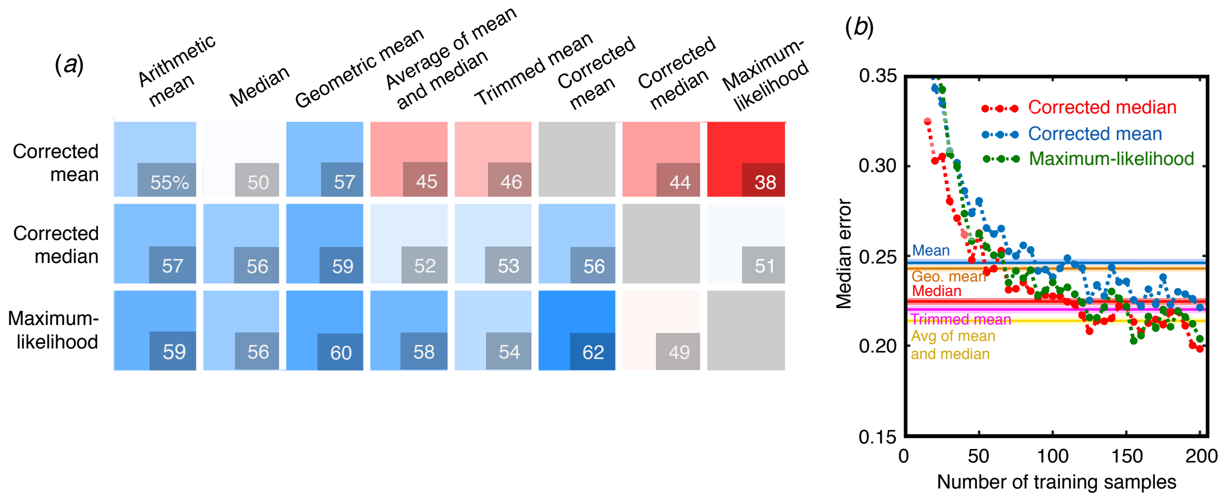


Figure S5: **The overall relative performance of the aggregation measures, using data drawn from the experimental datasets.** (a) The percentage of simulations in which the measure indicated in the row was more accurate than the measure indicated in the column. The three new measures are listed in the rows and are compared to all eight measures in the columns. Colors correlate with percentages (blue: >50%, red: <50%). (b) The median error of the three new aggregation measures (corrected median, dashed red line; corrected mean, dashed blue line; maximum-likelihood measure, dashed green line) as a function of the size of the training dataset. The three new aggregation measures are compared against the arithmetic mean (solid blue), median (solid red), the geometric mean (orange), the average of the mean and the median (yellow), and the trimmed mean (magenta). The 95% confidence interval are displayed for the latter measures, which are not a function of the size of the training dataset.

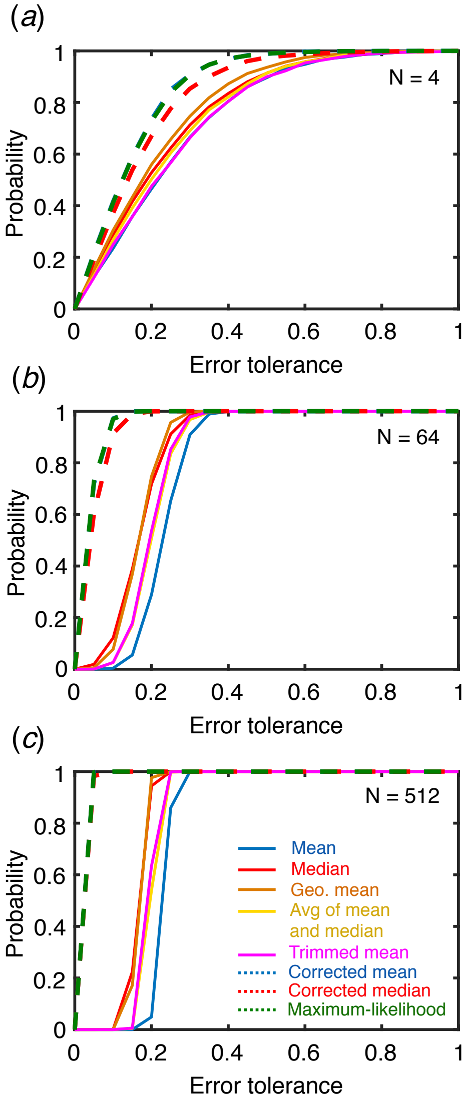


Figure S6: **The effect of the test dataset size and error tolerance level on the relative accuracy of the aggregation measures for numerosity** $\ln(J) = 4$ $(J = 55)$**, using simulated data.** The probability that an aggregation measure exhibits a relative error (defined as $|X - J|/J$, where $X$ is the value of an aggregation measure) less than a given error tolerance, for test dataset size (a) 4, (b) 64, and (c) 512. In panel (a), the lines for the corrected mean and the maximum-likelihood measure, and the lines for the the arithmetic mean and trimmed mean, are nearly identical; in panel (b), the lines for the corrected mean and maximum-likelihood measure, and the lines for the average of the mean and median and the trimmed mean, are nearly identical; and in panel (c), the lines for the corrected mean, corrected median, and the maximum-likelihood measure, the lines for the median and the geometric mean, and the lines for the average of the mean and median and the trimmed mean, are nearly identical.

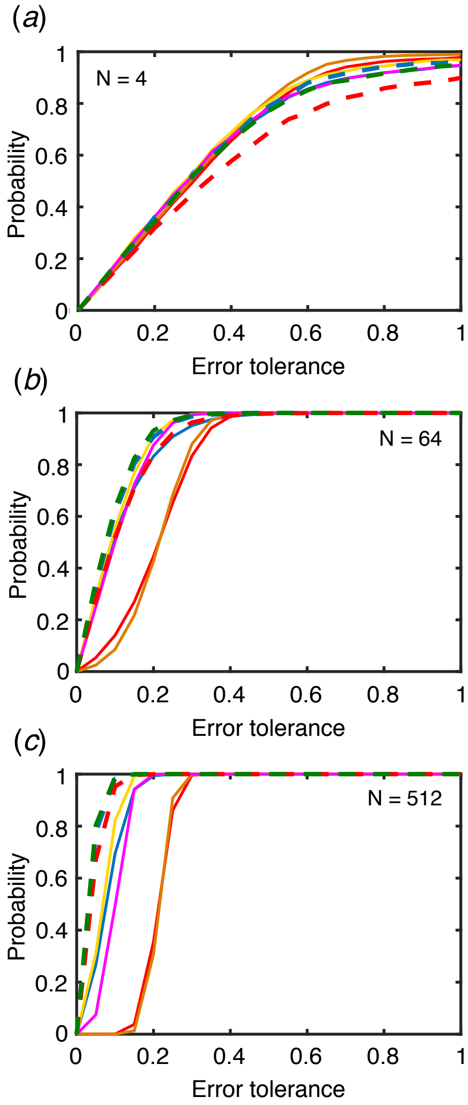


Figure S7: **The effect of the test dataset size and error tolerance level on the relative accuracy of the aggregation measures for numerosity** $\ln(J) = 7$ $(J = 1097)$**, using simulated data.** The probability that an aggregation measure exhibits a relative error (defined as $|X - J|/J$, where $X$ is the value of an aggregation measure) less than a given error tolerance, for test dataset size (a) 4, (b) 64, and (c) 512. Line colors are same as in Figure S6. In panel (a), the lines for the arithmetic mean and the trimmed mean, and the lines for the median and the average of the mean and median, are nearly identical; in panel (b), the lines for the corrected mean and maximum-likelihood measure are nearly identical; and in panel (c), the lines for the corrected mean and the maximum-likelihood measure are nearly identical.

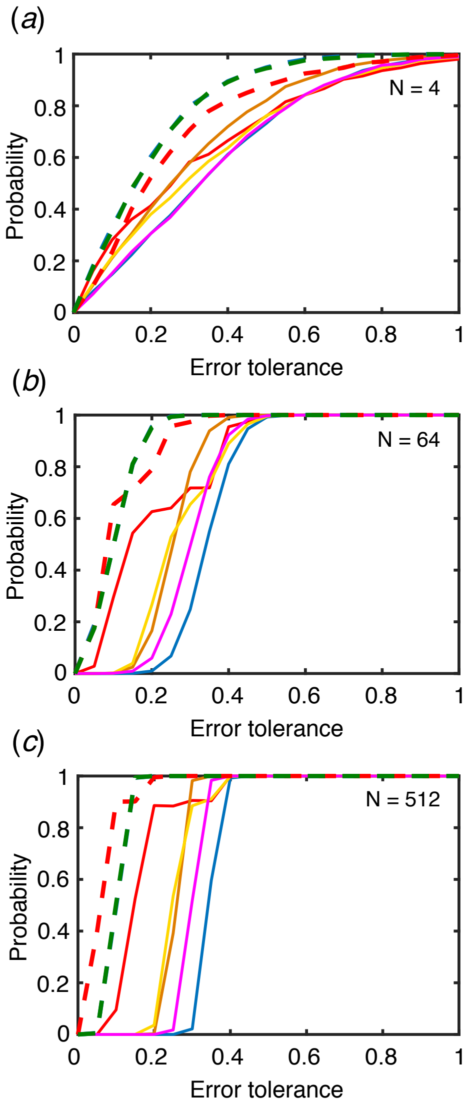


Figure S8: **The effect of the test dataset size and error tolerance level on the relative accuracy of the aggregation measures for numerosity $J = 54$, using data drawn from the experimental datasets.** The probability that an aggregation measure exhibits a relative error (defined as $|X - J|/J$, where $X$ is the value of an aggregation measure) less than a given error tolerance, for test dataset size (a) 4, (b) 64, and (c) 512. Line colors are same as in Figure S6. In panel (a), the lines for the corrected mean and the maximum-likelihood measure, and the lines for the arithmetic mean and the trimmed mean, are nearly identical; in panel (b), the lines for the corrected mean and maximum-likelihood are nearly identical; and in panel (c), the lines for the corrected mean and the maximum-likelihood are nearly identical.

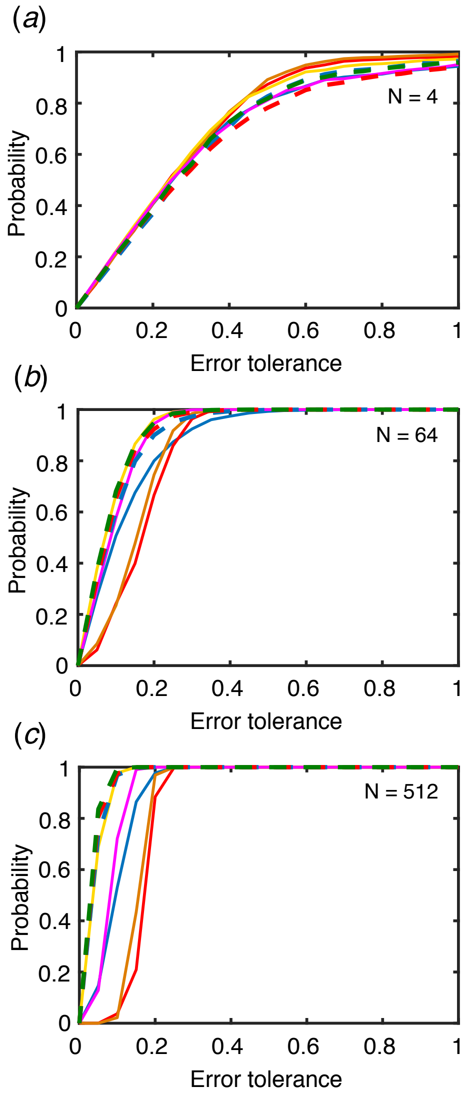


Figure S9: **The effect of the test dataset size and error tolerance level on the relative accuracy of the aggregation measures for numerosity $J = 659$, using data drawn from the experimental datasets.** The probability that an aggregation measure exhibits a relative error (defined as $|X - J|/J$, where $X$ is the value of an aggregation measure) less than a given error tolerance, for test dataset size (a) 4, (b) 64, and (c) 512. Line colors are same as in Figure S6. In panel (a), the lines for the corrected mean and the maximum-likelihood measure, and the lines for the arithmetic mean and the trimmed mean, are nearly identical; and in panel (c), the lines for the corrected mean, the corrected median, and the maximum-likelihood are nearly identical.

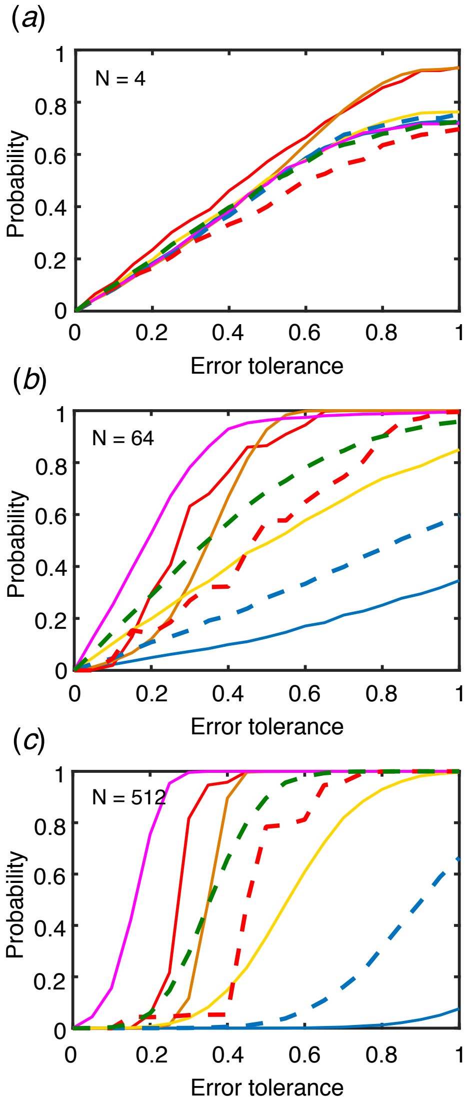


Figure S10: **The effect of the test dataset size and error tolerance level on the relative accuracy of the aggregation measures for numerosity $J = 27852$, using data drawn from the experimental datasets.** The probability that an aggregation measure exhibits a relative error (defined as $|X - J|/J$, where $X$ is the value of an aggregation measure) less than a given error tolerance, for test dataset size (a) 4, (b) 64, and (c) 512. Line colors are same as in Figure S6. In panel (a), the lines for the arithmetic mean and the trimmed mean are nearly identical.

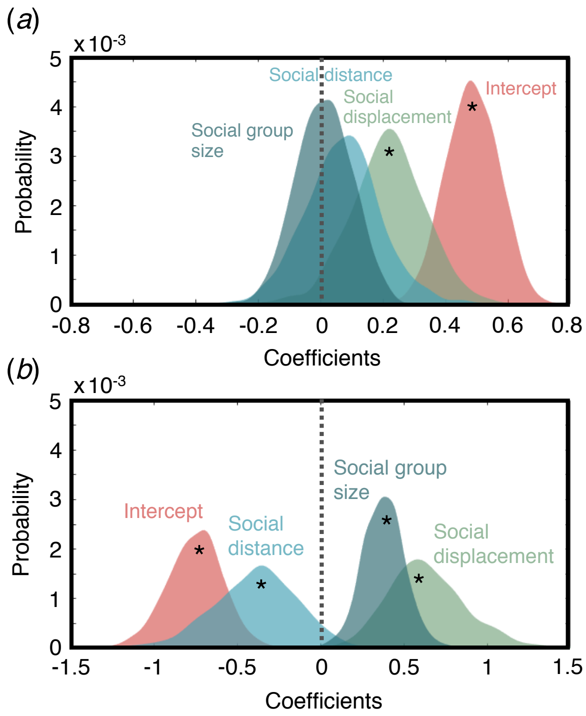


Figure S11: **The posterior distributions of the coefficients for the social displacement, social distance, social group size, and the intercept**, for (a) the model describing the probability that an individual chooses to use, or ignore, social information and (b) the model describing the strength of the social influence weight $\alpha$, for those individuals who chose to use social information. Note that the intercept for both models sets a baseline propensity to utilize social information (either probability or social influence weight) in the absence of other cues, where positive (negative) values indicate a increased (decreased) baseline propensity. Distributions with 95% credible intervals that do not include 0 are marked with an asterisk.

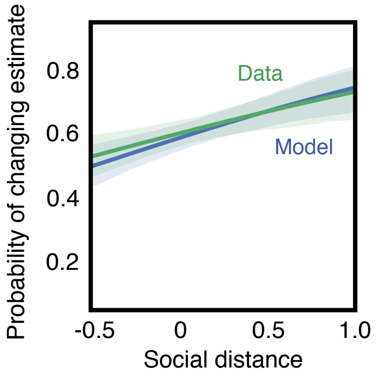


Figure S12: **Comparison of the posterior predicted probability of changing one's estimate in response to social information as a function of social distance (blue) with the observed probability from the data (green).** Shaded areas indicated 95% credible interval.

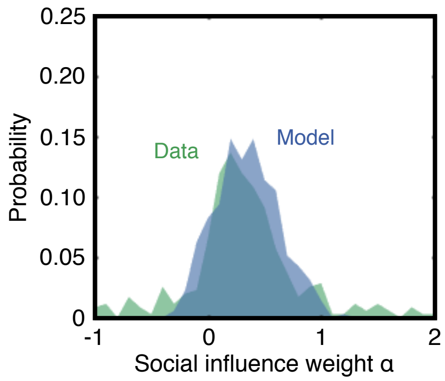


Figure S13: **Comparison of the posterior predicted distribution of social influence weights $\alpha$ (blue) with the observed distribution of $\alpha$ (green).**

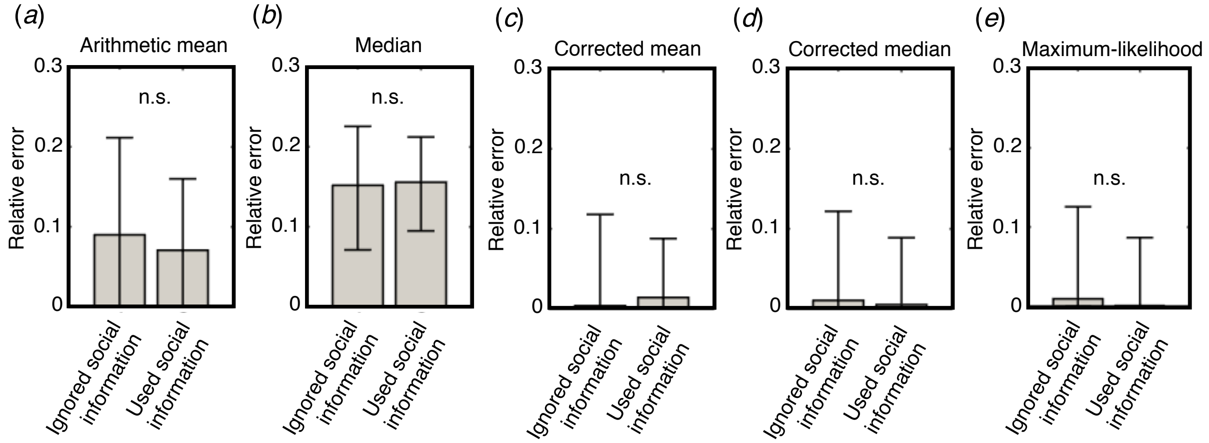


Figure S14: **Participants who ignored social information were not more accurate than those who utilized social information in their final estimates.** Shown are the maximum-likelihood relative error and 95% confidence intervals for (a) the arithmetic mean, (b) median, (c) corrected mean, (d) corrected median, and (e) maximum-likelihood measure. To generate confidence intervals, we identified all combinations of the parameters of the log-normal distribution which fell within the 95% confidence bounds. Then, we calculated and plotted the lowest and highest relative error for the values of the aggregation measures using those parameter combinations.
